# Empirically-Derived Synthetic Populations to Mitigate Small Sample Sizes

**DOI:** 10.1101/441238

**Authors:** Erin E. Fowler, Anders Berglund, Michael J. Schell, Thomas A. Sellers, Steven Eschrich, John Heine

## Abstract

Limited sample sizes can hinder biomedical research and lead to spurious findings. The objective of this work is to present a new method to generate synthetic populations (SPs) from sparse data samples to aid in modeling developments. Matched case-control data (n=180 pairs) defined the limited samples. Cases and controls were considered as two separate limited samples. Synthetic populations were generated for these observed samples using multivariate unconstrained bandwidth kernel density estimations. We included four continuous variables and one categorical variable for each individual. Bandwidth matrices were determined with Differential Evolution (DE) optimization driven by covariance comparisons. Four synthetic samples (n=180) were constructed from their respective SP for comparison purposes. Similarity between the observed samples with equally sized synthetic-samples was compared under the hypothesis that their sample distributions were the same. Distributions were compared with the maximum mean discrepancy (MMD) test statistic based on a Kernel Two-Sample Test. To evaluate similarity within a modeling context, Principal Component Analysis (PCA) score distributions and residuals were summarized with the distance to the model in X-space (DModX) as additional comparisons.

Four SPs were generated with the optimization procedure. The probability of selecting a replicate when randomly constructing synthetic samples with n=180 was infinitesimally small. The MMD tests indicated that the observed sample distributions were similar to the respective synthetic distributions. For both case and control samples, PCA scores and residuals did not deviate significantly when compared with their respective synthetic samples.

The reasonableness of this SP generation approach was demonstrated. This approach produced synthetic data at the patient level statistically similar to the observed samples, and thus could be used to generate larger-sized simulated data. The methodology coupled kernel density estimation with DE optimization and deployed novel similarity metrics derived from PCA. The use of large-sized synthetic samples may be a way to overcome sparse datasets. To further develop this approach into a research tool for model building purposes, additional evaluation with increased dimensionality is required; moreover, comparisons with other techniques such as bootstrapping and cross-validation will be required for a complete evaluation.

## 1. Background

Often data is limited in biomedical research, hindering scientific discovery. There are numerous reasons for sparse sample sizes, including low disease-incidence rates, rare diseases (1, 2), underserved/underrepresented subpopulations (3), the inability to share data across institutions due to privacy concerns (1), large-dimensionality relative to the number of patient samples [as the case in *omics* research] (4, 5), and limited timeframes for data collection, often due to funding cycles coupled with low disease incidence. These represent major barriers that hinder adoption of decision models in clinical practice.

Before applying a decision model in the clinical setting, potential models are typically explored during a discovery phase and then validated. The discovery phase includes analyzing samples from the respective populations to find measures (patient variables) related to the desired clinical endpoint, creating a model, and optimizing the model. Once the model parameters are fixed, the model can be evaluated with new data without further modeling adjustments. Both stages of development are dependent upon having access to adequately sized samples, which is often the major bottleneck. Inadequate sample sizes can lead to either false positive discoveries due to overfitting (6) or to false negative results due to reduced power. Automated variable selection, pretesting potential predictor variables, and dichotomizing continuous variables are all contributing factors to spurious findings (7). The extent that triaged variables and analyses are discussed in the literature is limited. Harrell (8) proposed a rough rule of thumb to decrease the risk of overfitting when modeling data. Briefly for binary data, the number of variables (more aptly degrees of freedom) considered when model building should be at most L/10 range, where L is the sample size of the smaller group, and it includes all variables examined even if they did not make it into the specified model (9). Others have suggested a lower threshold of L/5 may be sufficient (10). It is our premise that it is better to induce model failure during the discovery phase rather than with more costly later studies. It may not always be feasible to collect sufficient data during the initial development stages to thoroughly evaluate a given model. Methods to increase sample sizes synthetically, as proposed, have the potential to contribute significantly in these situations.

There are several techniques currently available to address sparse sample conditions. Bootstrapping (11) is a highly successful statistical resampling method often used when access to multiple samples is limited, but the method has noted deficiencies when applied to small sample sizes (12). A major concern is that a bootstrap sample only uses values seen in the sample. Other resampling techniques include various forms of cross-validation. Leave-one-out cross-validation produces a stringent test that underestimates the performance but has a drawback; in some instances removing one sample from the dataset will not produce enough variation (6). More generally, cross-validation can be highly variable (13). Cross-validation depends on repeatedly re-training to estimate the error rates. As such, it does not result in a specific set of model parameters, but rather parameters for each step of cross-validation. Another technique includes splitting the sample into training and validation subsets. Splitting the sample provides for independent replication but is dependent upon a dataset that is large enough to separate. As a potential alternative approach to resampling techniques, we will borrow from synthetic population (SP) generation methods (14) and explore generating synthetic patient data.

Synthetic population generation methods are frequently used in policy planning studies (15), such as land use and transportation research (16–18) when accessing data may be prohibitive due to privacy or cost issues (16). The objective is to construct the joint probability distribution for the population of interest (target) given the aggregated data (marginal distributions or tabulated data) and a sample of disaggregated data (joint distribution) with specified constraints. Established approaches (14, 15, 19) include deterministic re-weighting techniques and methods that use some form of stochastic component in the synthesis. Iterative proportional fitting (IPF) is a deterministic re-weighting method that is a well-tried approach (20) based on iteratively adjusting contingency tables relative to the given constraints as detailed by Lomax and Norman (21). Although IPF extends to any number of dimensions (22), there are noted efficiency problems for large dimensionality (23). The limitations with IPF are discussed in detail elsewhere (18, 21, 23), which include having too many zero cells, which may hinder convergence, and problems with high-dimensionality. The conditional probability approach uses the observed sample to estimate the target population by randomly sampling one variable at a time. This approach becomes cumbersome for a larger number of variables, and the outcome is dependent upon how the constraints are introduced (i.e. ordered) into the stochastic distribution reconstruction (19). Combinatorial optimization (CO) techniques are stochastic methods that include hill-climbing and simulated annealing (19, 24). In CO approaches, the observed sample is often assumed to be large compared to the target population. Random samples are drawn from the larger population and compared with the target population constraints; the element that enhances the goodness-of-fit becomes part of the target population (19, 24, 25). Hill-climbing has the advantage of speed (24) but can become trapped in suboptimal solutions (19, 24). Simulated annealing is a refinement of hill-climbing that permits the possibility of backing out of a suboptimal solution but, as a consequence, may lose performance (19, 24).

In this report, a new methodology for generating SPs is presented with some initial findings to demonstrate proof-of-concept. In this section, a brief outline of our approach is provided, followed by an in-depth description in the Methods Section. We define an SP as a population of conceptually unlimited individuals that is inferred from an observed sample. We refer to the case and control groups as samples from distinct populations (i.e. case and control populations) with 180 individuals in each sample. Synthetic samples are defined as 180 synthetic individuals selected from respective case or control SP. The boundary conditions differ from some of the approaches discussed above. We start with an observed sample, which is generally sparse (with respect to the population), with the objective of constructing a SP from which synthetic samples can be constructed. Synthetic samples were constructed randomly from the SP and compared with the corresponding observed sample. Multivariate kernel density estimation with an unconstrained bandwidth matrix was used to create these SPs. Differential Evolution (DE) optimization (26) was used to determine the bandwidth parameters for the kernel density estimation by matching the covariance structure between the observed sample and the synthetic samples. Multiple forms of analyses were used to thoroughly evaluate the similarity between the observed sample and synthetic samples: (i) the maximum mean discrepancy (MMD) metric with the Kernel Two-Sample Test (27) was used to compare the distributions; (ii) covariance matrices were compared; and (iii) to evaluate this approach within a modeling context, novel similarity metrics derived from Principal Component Analysis (PCA) were employed to compare the multivariate structure.

## 2. Methods

### 2.1 Sample and Synthetic Data

Observed samples are from a matched case-control study (n = 180 pairs) comprised of women who underwent mammography at Moffitt Cancer Center. Cases and controls were considered as two populations and the observed samples (case-sample and control-sample) were treated as two samples from two different distributions. Women with first time unilateral breast cancer defined the case-sample. The control-sample included women without a history of breast cancer. Controls were individually matched to breast cancer cases by age (±2 years), and other factors not relevant to this work. The data accrual and nuances of the matching were discussed previously (28–31). Although this matched case-control data was constructed and investigated for epidemiologic purposes previously, we used it to explore this SP generation methodology in a more realistic situation, where the epidemiologic context is irrelevant. For feasibility, we selected these variables for each individual: (1) age measured in years [yr]; (2) height measured in inches [in], (3) mass [kg], (4) breast density measure (percentage), referred to as PD; and (5) menopausal status [MS]. Measures 1-4 are continuous variables rounded to integer values and measure 5, corresponding to MS, is binary with MS = 0 for menopausal women and MS = 1 otherwise; in the expressions below, λ was used for the respective MS index for brevity. We used the inch as the height unit for because larger metrics (meters or feet) were too coarse, whereas a smaller metric, such as cm, was too resolved.

Synthetic samples were generated similar to the way the observed samples were established. Because controls were matched to cases (not selected randomly), we generated two synthetic populations from which synthetic samples can be constructed. For a given synthetic population, 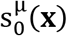, we considered the four integer variables as components of a vector referred to as, **x**, and generated the conditional population based on MS: 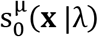. In this expression the index, μ, defines the case (μ=1) and control (μ=0) status, and the subscript, 0, indicates the population before scaling and normalization were applied. The general expression for both populations can be expressed explicitly in one given by

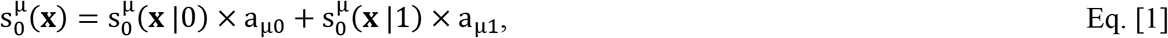

where a_μλ_ are the respective weights derived for the corresponding samples: a_00_ = 0.773 and a_01_ = 1.0 − a_00_, derived from the control-sample, and a_10_ = 0.788 and a_11_ = 1.0 − a_10_, derived from the control-sample. A synthetic sample was produced by drawing a realization at random from the respective synthetic population using the methods discussed below. We note in the modeling situation, a given synthetic-control realization must be matched to its case. This can be achieved by drawing a random realization from a restricted region of 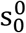 corresponding to the matching variables. In this report, synthetic case and control samples were generated and evaluated separately forgoing the matching necessity. As indicated in Eq. [1], there are two subsamples for each observed sample based on MS and two corresponding subpopulations. The number of individuals for the respective subsample is defined by N=180×a_μλ_. These subsamples were used to generate their respective subpopulations, referred to as components below.

### 2.2 Synthetic Population Density Estimation

To generate synthetic populations, we used a multivariate kernel density to estimate the underlying synthetic joint density derived from each sample. We defined an m component row vector for the **i**^th^ individual (an element from a given sample) **x**_i_ =(x_1_,x_2_,x_3_…x_m_) = [x_1_,x_2_,x_3_…x_m_]^T^. In our work m = 4. The components of **x**_i_ are x_ij_ and are sometimes referred to as features or predictor variables. All variables were rounded to integer values to reduce the computations required for the kernel density estimation. We defined the prospective m component column vector as **x** with components as x_j_, which is a synthetic entity. A given sample assumes the role of the training dataset for the respective SP generation. For motivational purposes, we first illustrate the simpler formulism. It may be shown (32) that the estimated joint density function for a sampled population with N individuals for the constrained bandwidth estimation is given by

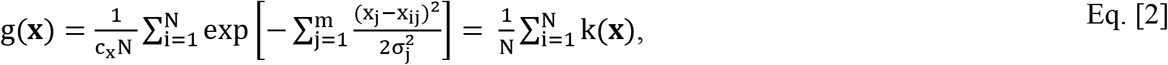

where we have assumed independent samples, c_x_ is a multivariate normalization constant, and σ_j_ are the bandwidth parameters. We used a Gaussian kernel, although there are many valid kernel choices. This operation is performed for each **x**. Equation [2] is too restrictive for our application because it lacks variable interaction. The above expression can be generalized to capture the relevant covariance structure (33) giving the unconstrained bandwidth expression

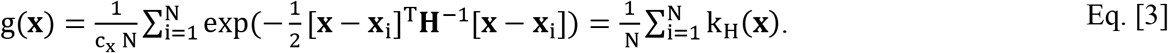

We have included the normalization constants in Eqs. [2–3] for completeness, although they were dropped in the analyses. **H** is the bandwidth matrix, which is symmetric and positive definite. These conditions on **H** ensure each term in the sum provides a valid probability density increment. If **H** is diagonal, the above expression reduces to Eq. [2]. Initial experimentation showed we could drop the ½ scale factor in the argument of Eq. [3] and obtain crude estimate of **H** as the covariance. Using Eq. [3], the population form that includes the subpopulations for both cases and controls is explicitly expressed as

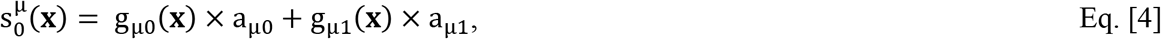

corresponding to Eq. [1]. Thus, the optimization problem requires two estimations per synthetic population due to the MS splitting.

### 2.3 Bandwidth Matrix Determination

The bandwidth parameters affect both the degree of smoothing and orientation. Selecting the optimal bandwidth parameters for unconstrained **H** is critical and non-trivial. In particular, bandwidth selection for the unconstrained multivariate problem is the subject of ongoing research, (34, 35). DE was used as the optimization method based on covariance comparisons for each subpopulation. The following optimization was used to construct each subpopulation separately from the respective subsamples. The N × m design matrix for a given subsample was defined as **X**_m_ with elements x_ij_, where the rows of **X**_m_ contain 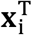. Mean centering the columns of **X**_m_ gives **X**. The covariance for a given subsample is expressed as

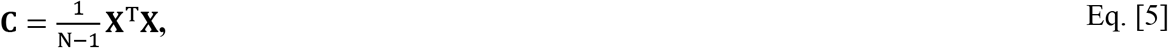

where **C** is an m×m matrix and N is the number of individuals in a given subsample. The covariance of a random sample of N individuals drawn from a given subpopulation is defined as **C**^syn^, using the same formulism expressed in Eq. [5]. For the optimization procedure, we used the normalized absolute difference matrix **E**, with elements: 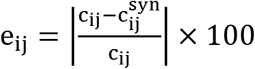, where **E** is m×m matrix. **H** is specified by minimizing the difference

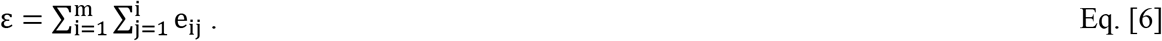

A brief description of DE is provided, aided by the schema shown in Figure 1 and Figure 2. Where possible, we followed the notation established by its founders (26). As such, the variable labeling and indexing introduced for this development are restricted to this section unless noted. This is a global optimization technique that constructs a solution via competition. We start with a randomly initialized vector field (zeroth generation), where the components, indexed by j of a given vector correspond to the elements of the bandwidth matrix, **H**, defined previously. A uniform random variable (rv) was used to generate these components constrained to ±0.10×estimated respective covariance from a given subsample. This DE-population was comprised of Np vectors (i.e. the DE-population size) with D_n_ components, where g is the generation index. We refer to the entire DE-population as **w**_g_ and label its members as **w**_ig_, where i is the member index. For this development, D_n_ is the number of unique elements in **H**. For a given g, we cycle through i = 1 to Np (Np is defined below) and select **w**_ig_ at each increment as illustrated in Figure 1. A mutant vector is constructed by adding a weighted difference between two vectors to a third vector, all randomly selected from **w**_g_ expressed as

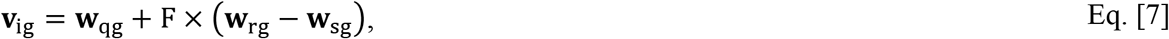

subject to this condition: q ≠ r ≠ s ≠ i, where q, r, as s are uniformly distributed, [1, Np], integer rvs; this step induces diversity. The weight, F, is referred to as the amplification factor and controls the mutation size or effect. The mutant vector is used to construct a trial vector defined as **u**_ig_ that competes with **w**_ig_ shown in Figure 1. A trial vector is constructed by permitting the possibility (at random) of crossing components of this mutant vector with the components of the target vector dictated by a crossover rate (CR) shown in Figure 2. The crossover operation occurs at the vector component level and dictates the mutation rate. This is to enhance the possibility that the trial vector has at least one component from the mutation (Figure 2). There are two avenues for the mutation to occur at each component: (i) if CR < a uniformly distributed, [0,1], rv or (ii) if a uniformly distributed [1,D_n_] integer rv = j. These random quantities are defined as c_j_ and d_j_, respectively. A given **u**_ig_ represents a mutation (most of the time) of **w**_ig_ derived from Eq. [7]. A given set of target and trial vectors compete to move the next generation by comparing their respective ε calculated with Eq. [6] after generating an SP for each vector. The vector that produces the smaller ε moves to the next to generation; Np comparisons of **w**_ig_ with **u**_ig_ produce the next generation population defined as **w**_g+1_. Thus, successive generations of **w**_g_ become incrementally more fit (i.e. decreasing ε). This process is performed generation by generation (Figure 1) until convergence. Although **H** has m^2^ components, we force symmetry on the solution and used m×(m+1)/2 components; thus all the vectors in the DE application have D_n_ = m×(m+1)/2 components. For this DE-population size, we use Np = 10×D_n_= 10×10=100. We also force the positive definite property on the solution by checking **H** for positive Eigenvalues. When a possible solution is not positive definite we assign a large error term to the competition rather than using Eq. [6]. In the event that both vectors produce a non-positive definite **H**, the target vector moves to the next generation. Thus, as the generations unfold, the possibility of forming a non-positive definite potential solution tends to zero.

**Figure 1:**
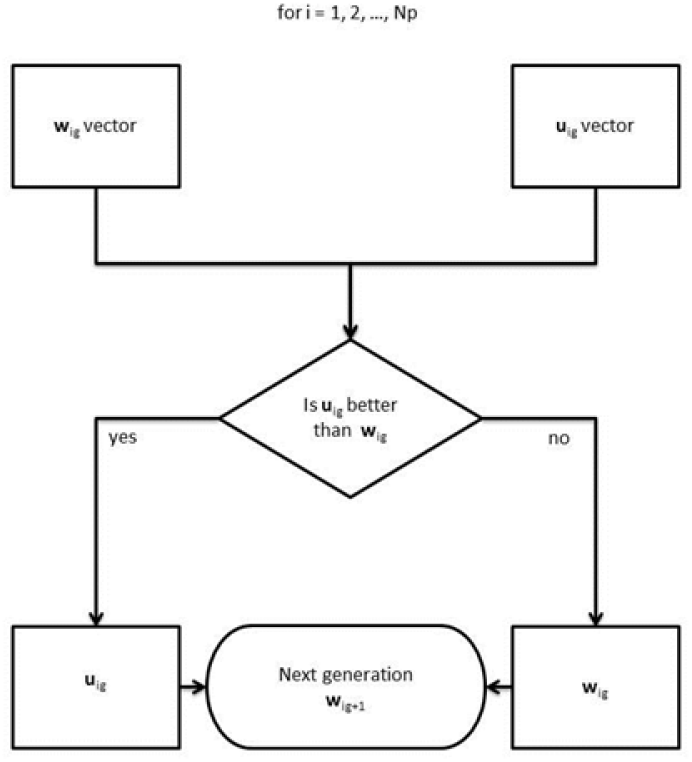
Differential Evolution Trial Competition: This illustrates the basic competition for one trial within a given generation. There are Np trial competitions per generation. The **i**^th^ vector from the current generation, **w**_ig_, competes with a trial vector, **u**_ig_. The trial vector is a mutation of **w**_ig_ with attributes derived from the population **w**_g_ shown in Figure 2.

**Figure 2:**
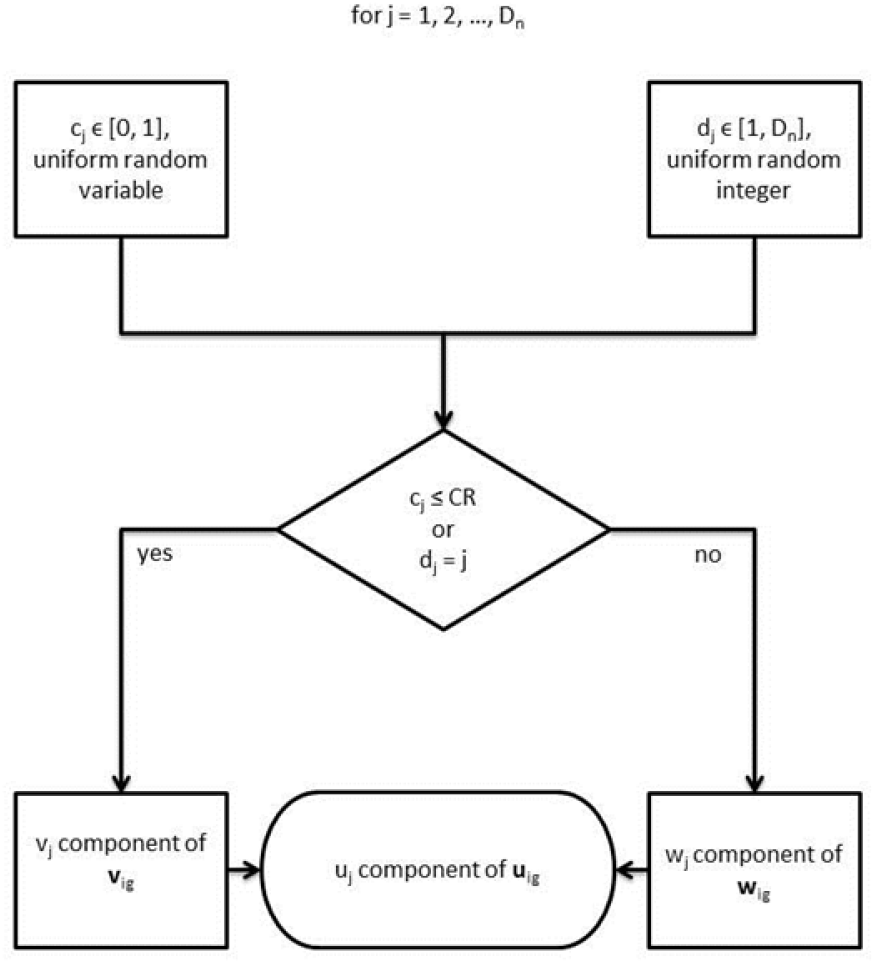
Trial Vector Construction: The trial vector, **u**_ig_, is constructed with components from **w**_ig_ and the mutant vector **v**_ig_. The **v**_ig_ construction is shown in Eq. [7]. The Vectors **u**_ig_ and **w**_ig_ compete shown in Figure 1.

### 2.4 Synthetic Sample Construction

Synthetic samples of N individuals were constructed during the optimization procedure corresponding to the four subpopulations. We adapted the standard method of mapping a uniformly distributed random variable to a random variable with a specified (target) probability distribution function (pdf), sometimes referred to as inverse sampling. For completeness, we outlined this standard transform method to put our approach in context. We define two random variables: (i) u, over this range [0,1] with a pdf given by p_u_(u) = 1, i.e. u is uniformly distributed random variable; and (ii) t over this range (−∞, ∞) with the target pdf labeled as p_t_(t), where p_t_(t) is arbitrary. We then determined the function q(u) = t, such that t is distributed as p_t_(t). The solution conserves probability using the cumulative functions

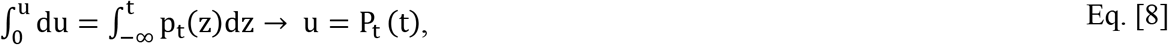

where P_t_(t) is the cumulative probability function for t. In passing, the above shows that mapping any random variable with its own cumulative function produces a uniform random variable. Although we used Eq. [8], we show that standard and equivalent solution expressed as

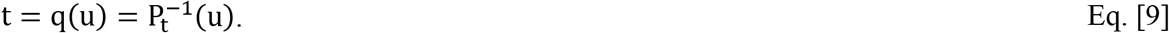

To construct a given SP, we first scaled its g(**x**) components defined in Eq. [3] with reasonable approximations. For simplicity, the case-control and MS indices have been suppressed. We assumed that there were approximately W women in each subpopulation (W ~ million) and scaled g(**x**): h_1_(**x**) = W× g(**x**). For all **x** meeting the condition h_1_(**x**) < 1, we set h_1_(**x**) = 0, which gives h(**x**). This normalization permits only whole synthetic individuals. Incorporating the indexing, the final expression for both SPs is given by

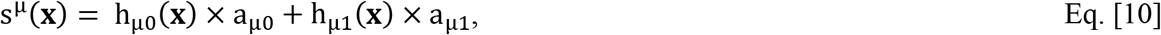

which follows from Eq.[4].

As an alternative to adding the components, Eq. [10] was realized stochastically in our experiments. To construct a given SP, a component was first selected at random dependent upon a_μλ_ to comply with Eq. [10]. To sample a given component, we let u_1_ be a uniformly distributed, [1,w], integer rv, where w is the *area* under the specific h_μλ_(**x**) component, and is equivalent to the number of women in the component after the scaling and normalization. We assembled all of the unique combinations of the coordinates **x**= (x_1_,…,x_m_) into a one dimensional variable, z_i_, where z_1_ is combination 1, z_2_ is combination 2, z_i_ is the **i**^th^ combination, and continue the ordering to the last combination labeled as z_R_, where R is the number of unique combinations (after scaling and normalizing). We ordered the corresponding number of individuals with each combination into another one-dimensional variable, f_i_, also with R elements. It is important to recognize that combination ordering and labels are not important as long as the correspondence between z_i_ and f_i_ is maintained, which can be expressed as f_i_ = f_i_(z_i_). The cumulative function for f_i_ is given by

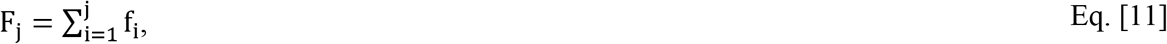

for j = 1 to R (i.e., F_R_ = w). This is the discrete and scaled analog of P_t_(t) expressed in Eq. [8]. A given individual was selected by generating a realization of a uniformly distributed rv, u_1_ defined above, and determining F_j−1_ < u_1_ ≤ F_j_. The interval F_j_-F_j−1_ gives f_j_ noting that f_j_ points to the **x** combination indexed by z_j_ producing one synthetic individual. We performed this operation repeatedly (i.e. selecting the h_μλ_(**x**) component and then selecting the individual with u_1_) to generate a sample of n individuals from a given SP. This solution is analogous to solving for a particular value of t (equating with **x**) given one realization of u expressed in Eq. [9].

### 2.5 Evaluation Methods

Multiple methods were used to compare the observed sample with synthetic samples when the optimization was completed. The Kernel Two-Sample Test (two-sample test) based on the MMD metric developed by Gretton et al (27) was used to compare the respective distributions. Additionally, the covariance structure was compared. To evaluate the similarity within the modelling context, two PCA-based methods were employed. In all comparisons, four synthetic samples were randomly constructed from the respective SP and compared with their respective observed sample.

A brief description of MMD metric and related test is provided using the same notation as Gretton et al (27), where possible. The comparison is based on a distribution free test to determine if two distributions are the same, noting it was designed to accept the null hypothesis. The MMD metric has multiple forms. When both distributions have the same number of elements (n), the squared MMD metric takes the following form used in this report

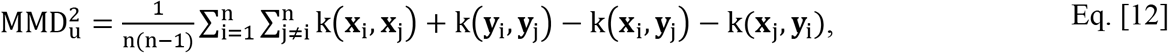

where subscript u indicates an unbiased estimate, **x**_i_ are the vectors for the individuals from a given sample and **y**_j_ are the vectors for the individuals from the respective synthetic sample, and the kernel terms refer to the forms expressed in Eqs. [2–3]. The null hypothesis test at significance level α (i.e. the distributions are the same) has the acceptance region expressed as

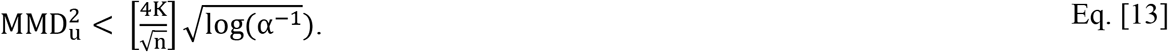

Equation [11] is referred to as the test-threshold below generated with α =0.05. The Type II error probability of this test approaches zero as n becomes large (27). The parameter K is determined from the bounds of the respective sample derived from the following inequality

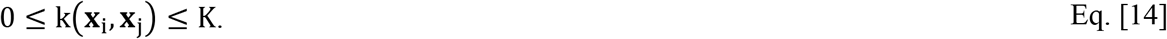

This test was performed twice for each observed sample and synthetic sample comparison; the test was performed with the kernel, k, expressed in Eq. [2] and with the kernel, k_H_, expressed in Eq. [3]. Because both the acceptance region and sample bounds scale with K, the quantities in Eq. [12] and Eq. [14] were generated with the non-normalized forms of k and k_H_ to keep the quantities from becoming too small. Each observed sample was compared with its respective synthetic samples using Eq. [12] with Eqs. [13–14]. The bandwidth matrices were derived from the covariance of the respective samples calculated without considering MS for these comparisons; when using diagonal **H**, variances determined from the respective sample were used to estimate the diagonal terms.

Additional experiments were performed to complement the comparison analysis. Experiments were performed to evaluate the degree to which synthetic samples included individual replicates from its respective observed sample. This was implemented by creating 1000 synthetic control-samples with n =180 from its SP and counting replicates. Due to the way the optimization process was controlled, covariance matrices were also compared between the observed samples and synthetic samples by evaluating the related matrix elements with 95% confidence intervals (CIs) separated by MS. To generate CIs for the sample covariance elements, 1000 bootstrap trials were performed. The covariance matrices were compared with the corresponding realizations of the respective synthetic covariance matrices. For the synthetic sample covariance elements, CIs were estimated by evaluating 1000 synthetic samples (each with n individuals) constructed from the respective SP.

The other objective was to assess the similarity within a modeling context to ensure that the multivariate structure between the samples and respective synthetic samples was similar. In this situation, we used a given observed sample as the reference and examined the similarity between respective synthetic samples with this reference. We applied two unique forms of analysis based on principal component analyses (PCA). First, the synthetic-samples were compared to the respective sample using PCA modeling; a PCA model derived from the sample was applied to the respective synthetic sample and predicted scores (expansion coefficients) were compared. This was evaluated by comparing the distributions along each individual principal component. Strong outliers were observable in the score plots, whereas the detection of moderate outliers required another approach. In the second approach, the distance to the model in X-space (DModX) statistic was used to evaluate differences between the sample and synthetic samples, where the residuals were evaluated. In short, the residuals should not deviate significantly. We note the PCA expansion and residuals are orthogonal complements. The PCA model was calculated (trained) on the sample and the variables were expanded with square and interaction-terms of the original variables. Each variable was centered and scaled to unit variance. The derived PCA models were then applied to the respective synthetic sample to predict the scores and generate DModX quantities. Residuals between the observed sample and synthetic samples were compared with a t-test. PCA models were calculated using Evince (version 2.7.9, Prediktera AB, Umea, Sweden). Box Plots with Violin lines were generated in MATLAB (R2017b, MathWorks Inc. Natick MA, USA) using the GRAMM toolbox (36). The SP and DE algorithms were developed in the IDL version 8.6 (Exelis Visual Information Solutions, Boulder, Colorado) programming environment.

## 3. Results

### 3.1 Sample Sparsity and Synthetic Replications

The two observed samples (e.g. cases and controls) were sparse samplings of their respective populations. The sparse characteristic of the control-sample (n=180) is illustrated in Figure 3. In this example, various projections of the control-sample distribution onto the mass-PD plane are shown. The top-left pane shows the projection for age = 58yr and height = 64in. Moving down the columns of the Figure 3 panes, we expanded the projection to include the age range from 58-59yr (middle) and 58-60yr (bottom). Moving right across the rows of the Figure 3 panes, projections correspond for height ranges from 64-66in and 64-68in, respectively. The most populated pane (58-60yr, height 64-68in) is on the bottom right, which includes 15 samples or 8% of the control-sample.

**Figure 3:**
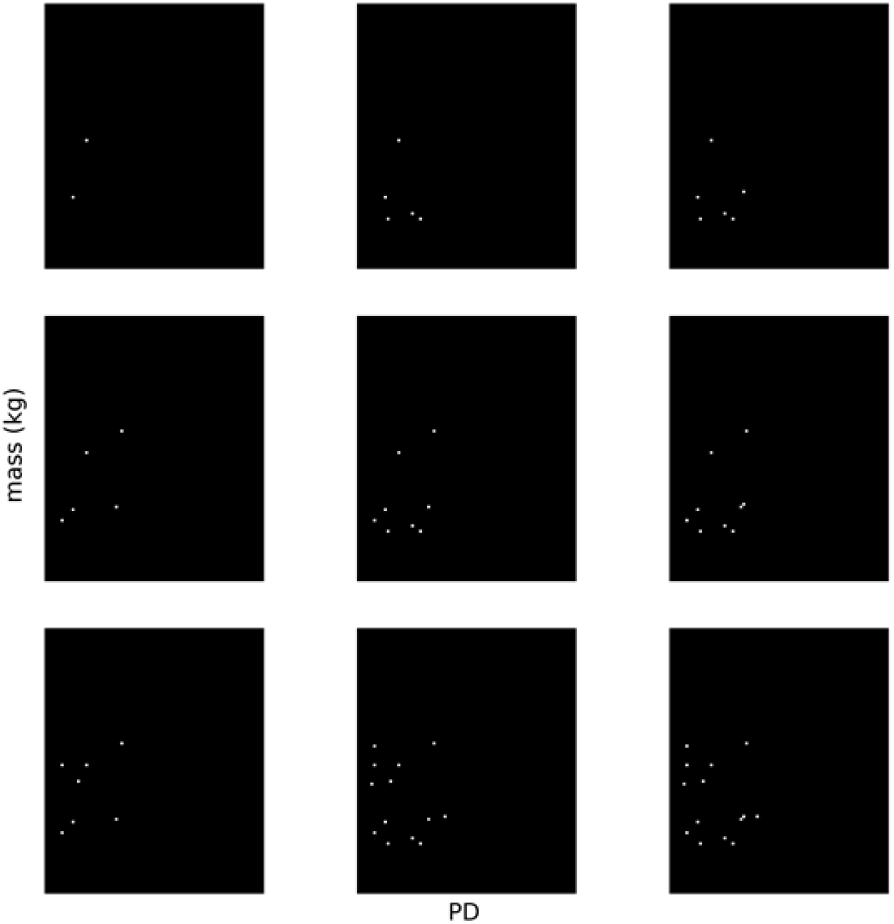
Control-sample sparsity illustration: This illustration shows multiple projections of the sample distribution onto the mass-PD plane. Mass (kg) is on the vertical axis and PD (breast density measure) on horizontal axis. The top left plot includes individuals with age = 58 years with height = 64in (most sparse plot in these examples). The top row (middle) includes individuals with a height range from 64-66in and top row (right) incudes individuals in 64-68in range. The second row shows the same height progression for women aged 58-59yr. Likewise, the third row (bottom) shows the same height progression for women aged 58-60yr (most populated in this illustration is on the bottom right). Each point represents one individual.

The SP generation methods produced a dense reconstruction of a population distribution, as illustrated in Figure 4. This shows a twodimensional slice through the four dimensional SP (left pane) at age = 58yr and height = 64in corresponding with the sparsest pane in Figure 3 (top left). Note the dense nature of this conditional distribution in comparison with Figure 3. Taking a profile along the horizontal direction at mass = 64kg, gives the associated conditional density (not normalized) for the PD (breast density metric) shown in the right pane (before thresholding). The smooth continuous curve illustrates the properties of the kernel reconstruction.

**Figure 4:**
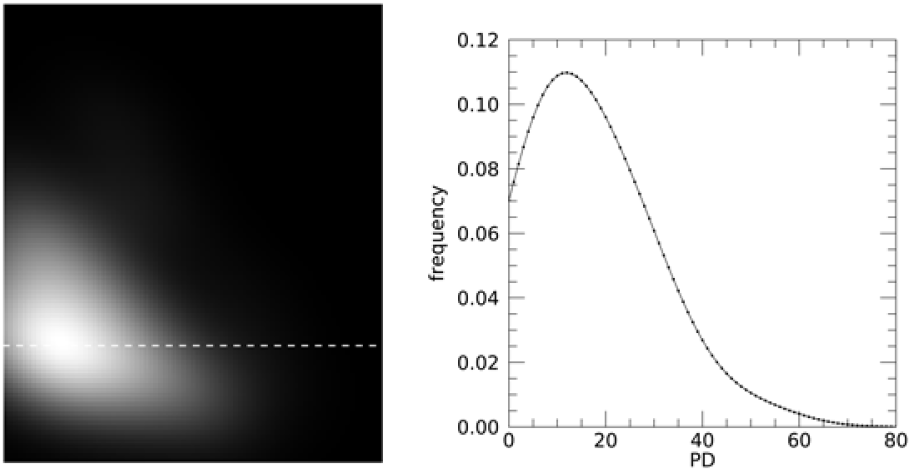
Synthetic control population reconstruction illustration. This shows a two-dimensional slice through the synthetic population (left pane) for women with age = 58yr and height = 64in. Mass (kg) is on the vertical axis and PD on the horizontal axis. This corresponds to the sparsest sample (n = 2) in Figure 3. The dashed line marks a profile for women with mass = 64kg. The corresponding distribution for PD (breast density measure) is shown on the right before integer rounding and normalization were applied.

We next applied the SP generation method to create 1,000 synthetic control-samples. We examined the frequency of generating a synthetic sample containing an individual that was present in the observed sample (replicate). The replicate evaluation revealed that the average number of synthetic samples (each with n= 180 individuals) was 18 (average) before a replicate individual from the observed sample was included. In 97.2% of the samples with replicates, there was only one replicate and in 2.8% of samples either 2-3 replicates were observed. In the similarity comparisons that follow, based on these findings, we conclude that the similarities do not arise from simple replicates of the observed sample.

### 3.2 Synthetic Population Generation Maintains Distribution Similarity

We compared four generated synthetic samples to the corresponding observed sample to determine if the distribution of individuals differed by using the two-sample kernel test. The findings from these two-sample tests using kH, the full bandwidth kernel, are shown in Table 1. In the comparisons between the control-sample with the four synthetic samples, the null hypothesis was accepted for each synthetic sample, indicating the synthetic samples have similar distributions to the observed sample. The findings comparing the case-sample with four synthetic samples were similar for k_H_ as well. Table 2 shows the tests based on k, the diagonal bandwidth kernel. The findings are similar to those shown in Table 1; in all comparisons, the null hypothesis was not rejected, suggesting that the SP generation method created synthetic samples that are distributed similarly to the observed sample. For reference, the bandwidth matrices determined with DE are provided in Table 3.

**Table 1.** Kernel Two-sample Test based on the full bandwidth kernel. Synthetic samples were compared to their respective case and control samples. Both K and the test threshold are provided. All quantities were generated with the full bandwidth kernel, k_H_ expressed in Eq. [3] with Eq. [12].

| Synthetic-sample | Control-sample<br>$MMD_u^2$<br>(K = 9.94E-01, threshold = 5.13E-0.1) | Case-sample<br>$MMD_u^2$<br>(K = 9.97E-01, threshold = 5.15E-01) |
| --- | --- | --- |
| 1 | -3.32E-03 | -4.45E-03 |
| 2 | -2.37E-03 | -1.96E-03 |
| 3 | -1.66E-03 | 2.52E-04 |
| 4 | 3.53E-05 | -9.03E-04 |

**Table 2.** A Kernel Two-sample Test based on the diagonal bandwidth kernel. Synthetic samples were compared to their respective case and control samples. Both K and the test threshold are provided. All quantities were generated with a diagonal bandwidth matrix kernel, k, expressed in Eq. [2] with Eq. [12].

| Synthetic sample | Control-sample<br>$MMD_u^2$<br>(K = 9.93E-01 threshold = 5.13E-0.1) | Case-sample<br>$MMD_u^2$<br>(K = 9.98E-01, threshold = 5.15E-01) |
| --- | --- | --- |
| 1 | -3.32E-03 | -3.87E-03 |
| 2 | -2.44E-03 | -1.93E-03 |
| 3 | -1.47E-03 | -2.57E-04 |
| 4 | -2.96E-04 | -1.10E-07 |

**Table 3.** Bandwidth Matrix Elements The bandwidth matrix, H, generated with DE is provided for each subgroup. These were used to generate the synthetic populations.

| Controls Menopausal n = 132 |  |  |  |  |
| --- | --- | --- | --- | --- |
|  | height (in) | Age (yr) | PD | Mass (kg) |
| Height (inch) | 5.16 | -2.08 | 2.93 | 5.20 |
| Age | -2.08 | 62.0 | -14.4 | -5.33 |
| PD | 2.93 | -14.4 | 185.6 | -48.1 |
| Weight (kg) | 5.20 | -5.33 | -48.1 | 141.3 |

| Cases Menopausal n = 142 |  |  |  |  |
| --- | --- | --- | --- | --- |
|  | height (in) | Age (yr) | PD | mass (kg) |
| Height (inch) | 6.27 | -4.94 | -2.28 | 10.6 |
| Age | -4.94 | 66.9 | -3.69 | -10.1 |
| PD | -2.28 | -3.69 | 188.3 | -59.4 |
| Weight (kg) | 10.6 | -10.1 | -59.4 | 164.8 |

| Controls non-Menopausal n = 48 |  |  |  |  |
| --- | --- | --- | --- | --- |
|  | height (in) | Age (yr) | PD | mass (kg) |
| Height (inch) | 6.68 | 2.14 | 11.9 | 2.23 |
| Age | 2.14 | 31.2 | -24.8 | -6.17 |
| PD | 11.9 | -24.8 | 242.0 | -65.3 |
| Weight (kg) | 2.23 | -6.17 | -65.3 | 73.3 |

| Cases non-Menopausal n = 38 |  |  |  |  |
| --- | --- | --- | --- | --- |
|  | height (in) | age (yr) | PD | mass (kg) |
| Height (inch) | 11 | -0.41 | -10.8 | 9.02 |
| Age | -0.41 | 19.7 | -10.4 | -2.85 |
| PD | -10.8 | -10.4 | 314.8 | -130.5 |
| Weight (kg) | 9.02 | -2.85 | -130.5 | 199.3 |

### 3.3. Covariance Comparisons

The covariance of a synthetic sample was compared to the corresponding observed sample. The covariance matrices for the control-sample and a synthetic control-sample are shown in Table 4 and Table 5 for the case-sample and a synthetic case-sample. Synthetic quantities did not deviate significantly from the respective sample quantities when considering intra-group pairwise comparisons.

**Table 4.** Covariance Matrices for Controls: These tables show the covariance matrix for the control-sample the left and a synthetic control-sample (one realization) on the right. Confidence intervals are provided parenthetically.

| Control-sample n = 180 |  |  |  |  |
| --- | --- | --- | --- | --- |
|  | height (in) | age (yr) | PD | mass (kg) |
| height (in) | 6.3<br>(5.1, 7.4) | -3.9<br>(-7.9, 0.0) | 6.6<br>(1.6, 11.6) | 3.9<br>(0.2, 7.7) |
| age | -3.9<br>(-7.9, 0.0) | 108.2<br>(88.7, 127.7) | -41.4<br>(-65.4, -17.4) | -2.8<br>(-16.6, 10.9) |
| PD | 6.6<br>(1.6, 11.6) | -41.4<br>(-65.4, -17.4) | 212.9<br>(167.3, 258.6) | -52.5<br>(-73.5, -31.5) |
| mass (kg) | 3.9<br>(0.2, 7.7) | -2.8<br>(-16.6, 10.9) | -52.5<br>(-73.5, -31.5) | 116.0<br>(84.1, 148.0) |

| Synthetic Control-sample n = 180 |  |  |  |  |
| --- | --- | --- | --- | --- |
|  | height (inch) | Age (yr) | PD | mass (kg) |
| height (inch) | 6.5<br>(5.4, 7.6) | -3.9<br>(-6.6, -1.3) | 11.4<br>(7.7, 15.1) | 6.3<br>(2.7, 9.8) |
| Age | -3.9<br>(-6.6, -1.3) | 105.8<br>(94.7, 116.9) | -40.5<br>(-52.8, -28.1) | -3.4<br>(-14.0, 7.2) |
| PD | 11.4<br>(7.7, 15.1) | -40.5<br>(-52.8, -28.1) | 185.0<br>(159.3, 210.8) | -35.9<br>(-53.3, -18.5) |
| mass (kg) | 6.3<br>(2.7, 9.8) | -3.4<br>(-14.0, 7.2) | -35.9<br>(-53.3, -18.5) | 128.4<br>(103.7, 153.0) |

**Table 5.**
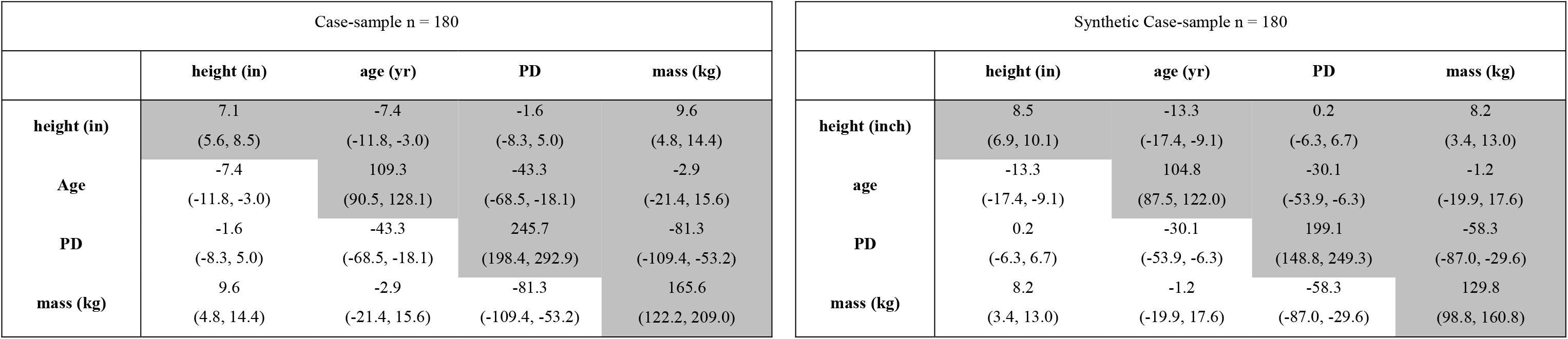
Covariance Matrices for the Cases: These tables show the covariance matrix for the case-sample on the left and synthetic case-sample (one realization) on the right. Confidence intervals are provided parenthetically.

### 3.4 PCA Comparisons

Characteristics of the synthetic-samples were also evaluated using PCA model comparisons with the corresponding observed sample. The associated findings are shown in Figure 5. Figure 5A (top left) shows the PCA scores for PC1 and PC2 (first two principal components) for the case-sample. The first principal component explains 26% of the variation, with the second explaining 18.2%, giving 44.2 % in total for this sample. These principal components are plotted together with the predictions of the four different synthetic-samples; these predictions were based on the trained PCA models for the case-sample and control-sample (two separate models were trained). We note, none of the synthetic samples contained replicates from the respective sample. The score plots show a distinct separation in all datasets due to MS (menopausal status). Approximately 99% of the individuals in the right group in Figure 3A had MS = 1 (about 1% had MS = 0), while the compact subgroup to the left consisted of individuals with MS = 0 (except one individual or approximately 0.14%). Interestingly, this artifact was a characteristic of the observed sample that is also present in the synthetic sample structure as well. The PCA score plot in Figure 3C shows similar behavior in that there are few differences between the observed sample and the synthetic-samples. Note, the separation due to MS was also captured. The second dimension (y-axis) is in the opposite direction for the control-sample compared to the case-sample. This is a common artifact in PCA because the signs are arbitrary. This is usually accounted for by multiplication of −1 (i.e. PCA is invariant under a sign change).

**Figure 5:**
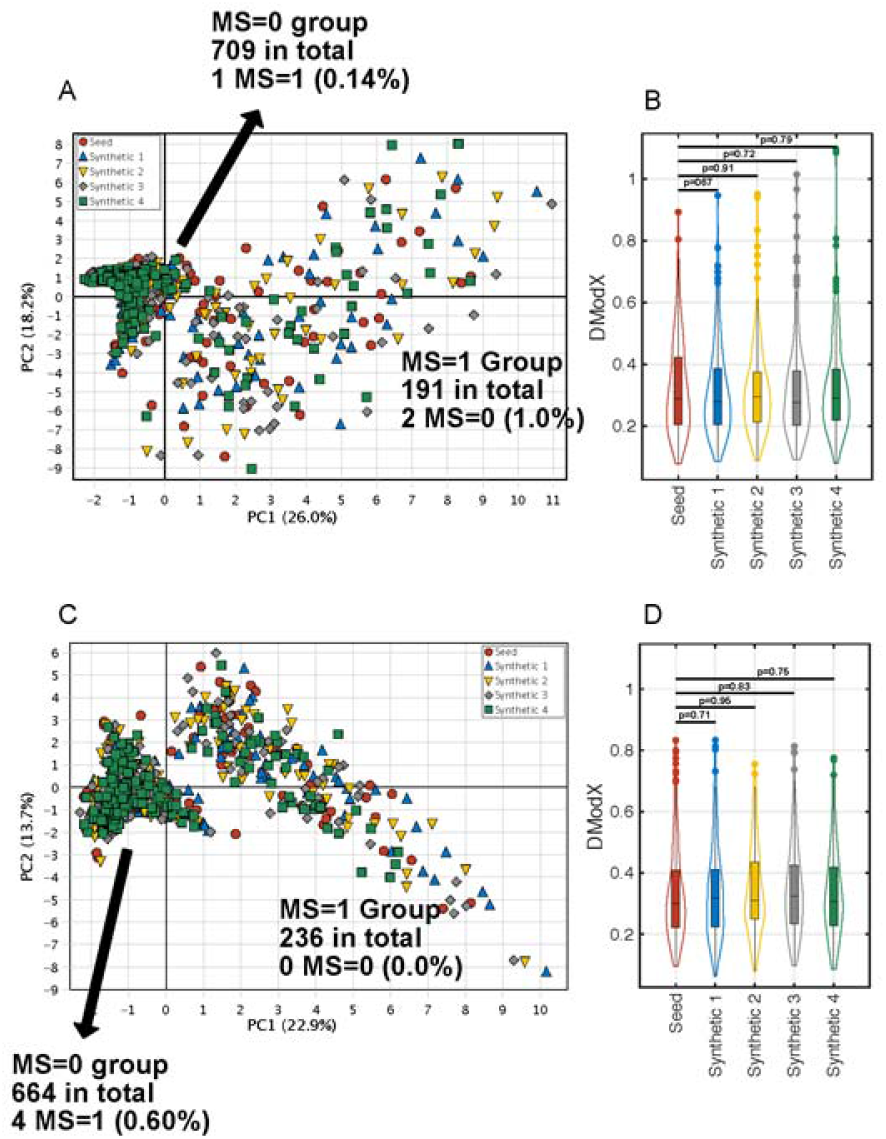
PCA models for the Observed and Synthetic Samples. The first two principal components derived from the case-sample are shown in **A** together with the predictions of the synthetic data. There is no visible difference between the observed and the four different synthetic datasets. The residuals, DmodX, are shown in **B** as a boxplot with violin density lines. The differences in the residuals between the observed and synthetic data were not significant. Similar results were noted for the control-sample in pane C and D.

Figure 5A illustrates that there is no visible deviation between the observed sample and the four synthetic samples, as represented by the PCA model elements PC1 and PC2. The variation not explained by the PCA model is shown in the DModX plot in Figure 5B for the case-sample. The DModX for each sample (observed sample and 4 synthetic samples) is represented by a boxplot and the distributions are shown as violin plots. In each plot, the p-value from the comparison with the sample residual is provided. The difference in each comparison was not significant. Because the DModX values across the 5 analyses were statistically similar, the sample and synthetic samples were essentially indistinguishable. The same analysis was also performed on the control-sample shown in Figure 5D, which produced similar findings.

## 4. Discussion

We presented a novel methodology to generate SPs given observed samples with limited size. The approach is based on an unconstrained multivariate kernel density estimation driven by an optimization procedure. The similarity of samples from the SP versus the observed sample was demonstrated with multiple evaluation methods. The two-sample tests indicated that the observed samples and equally-sized synthetic samples were similar for all comparisons. We also compared the covariance structure, which showed that the observed samples and synthetic samples did not deviate significantly. Within the modeling context, the feasibility of our approach was evident based on the PCA model comparisons. A benefit of the PCA approach is that all variables under consideration are evaluated simultaneously and repeatedly because each PCA dimension is a linear combination of the input variables. For either cases or controls, the respective synthetic-samples exhibited similar behavior under the PCA transform. We presented this approach to generate SP samples for model building purposes. The approach can also be used as a method of imputation for missing variables noting the entire dataset would be used in the estimation. The method also lends itself to generating sampling requirements for multivariate investigations given a population sample.

Although our study addressed many technical challenges, there are several limitations and nuances worth noting. We considered five variables. This dimensionality was sufficient to develop a general framework and provide initial feasibility. To make the approach into a model building tool, the dimensionality of the input space must be increased, necessitating more powerful data processing capabilities. Moreover, comparisons with accepted techniques such as cross validation and bootstrapping will be required to adequately rate the value of this approach. We considered four continuous and one dichotomous variable. Often, the input space could include a wider mix of continuous, ordinal, and nominal variables. We used a conditional probability approach to address the limited mixture of variables in this work. Additional methods of estimating the covariance between mixed variable types (37) are required to render the problem suitable for kernel processing and to reduce the number of conditional probability branches. Otherwise, extensive branching becomes a limitation for datasets with few samples. The process was driven by considering the covariance directly. It is not clear at this time if this is generally optimal in wider settings. Alternatively, the optimization could be driven by these PCA similarity metrics or other summary metrics such as the two-sample test used in this evaluation. Additional experimentation is required to determine the optimal method of driving the optimization process.

When this approach is more fully developed, it could produce synthetic-samples for model building exercises, providing the underlying data structure of the sample was captured in the synthetic population. Once a model is optimized, it can be initially validated with the sample population producing a fully specified model suitable for independent validation. In this report, we presented our methodology and demonstrated its feasibility. Because the technique is not yet fully developed, we do not provide comparisons with other techniques such as bootstrapping or cross-validation at this time. To make these comparisons relevant, our methodology will require scaling to accommodate higher dimensionality, which is planned future work.

## 5. Conclusions

The work presented in this report contributes to both kernel density estimation and synthetic population research. The optimization feedback loop can be modified to incorporate other endpoints. The work also introduced two related methods of analyzing structure similarity with PCA. These similarity metrics have applications well beyond those presented in this report. The use of synthetic samples may be useful to mitigate limited sample sizes for initial model building exercises providing our technique is evaluated under more general conditions (higher dimensionality and more variable types). Planned research includes addressing the limitations addressed above.

## Declarations

### Ethics and consent to participate

This study was approved by the Institutional Review Board (IRB), University of South Florida, Tampa, FL (IRB# Ame13_104715)

### Consent for publication

The work does not contain personal identifiers.

### Availability of data

The datasets used and/or analyzed during the current study are available from the corresponding author on reasonable request.

### Author contributions

EEF is a coauthor (co-first author), developed the computer code for the kernel mapping and differential evolution, and assisted JJH in the methods development; AB is a coauthor (co-first author) and developed the similarity metrics based on principal component analysis; MJS is a co-author and provided statistical expertise; TAS is a coauthor and provided epidemiologic expertise; SE is a coauthor and assisted in the methods development; JJH is the senior author, conceived of the plan and methods for synthetic data generation and developed computer code.

## References

1. Mascalzoni D, Paradiso A, Hansson M. Rare disease research: Breaking the privacy barrier. Appl Transl Genom. 2014;3(2):23–9. PubMed PMID: Medline:27275410. English.

2. Darquy S, Moutel G, Lapointe A-S, D’Audiffret D, Champagnat J, Guerroui S, et al. Patient/family views on data sharing in rare diseases: study in the European LeukoTreat project. Eur J Hum Genet. 2016;24(3):338–43. PubMed PMID: Medline:26081642. English.

3. Erves JC, Mayo-Gamble TL, Malin-Fair A, Boyer A, Joosten Y, Vaughn YC, et al. Needs, Priorities, and Recommendations for Engaging Underrepresented Populations in Clinical Research: A Community Perspective. J Community Health. 2017;42(3):472–80. PubMed PMID: Medline:27812847. English.

4. Lay Jr JO, Borgmann S, Liyanage R, Wilkins CL. Problems with the “omics”. Trends in Analytical Chemistry. 2006;25(11).

5. Micheel CM, Nass SJ, Omenn GS. Evolution of Translational Omics:: Lessons Learned and the Path Forward: National Academies Press; 2012.

6. Hawkins DM. The problem of overfitting. Journal of chemical information and computer sciences. 2004;44(1):1–12.

7. Babyak MA. What you see may not be what you get: a brief, nontechnical introduction to overfitting in regression-type models. Psychosomatic medicine. 2004;66(3):411–21.

8. Harrell FE, Jr., Lee KL, Mark DB. Multivariable prognostic models: issues in developing models, evaluating assumptions and adequacy, and measuring and reducing errors. Statistics in medicine. 1996 Feb 28;15(4):361–87. PubMed PMID: 8668867.

9. Harrell Jr FE. Regression Modeling and Validation Strategies 1997 [updated June, 1997]. Available from: http://biostat.mc.vanderbilt.edu/wiki/pub/Main/ClinStat/model.pdf.

10. Vittinghoff E, McCulloch CE. Relaxing the rule of ten events per variable in logistic and Cox regression. American journal of epidemiology. 2007;165(6):710–8.

11. Efron B, Gong G. A leisurely look at the bootstrap, the jackknife, and cross-validation. The American Statistician. 1983;37(1):36–48.

12. Isaksson A, Wallman M, Göransson H, Gustafsson MG. Cross-validation and bootstrapping are unreliable in small sample classification. Pattern Recognition Letters. 2008;29(14):1960–5.

13. Efron B, Tibshirani R. Improvements on cross-validation: the 632+ bootstrap method. Journal of the American Statistical Association. 1997;92(438):548–60.

14. Heppenstall A, Harland K, Smith D, Birkin M. Creating realistic synthetic populations at varying spatial scales: a comparative critique of population synthesis techniques. Geocomputation 2011 Conference Proceedings, UCL, London 2011. p. 1–8.

15. Smith DM, Clarke GP, Harland K. Improving the synthetic data generation process in spatial microsimulation models. Environment and Planning A. 2009;41(5):1251–68.

16. Barthelemy J, Toint PL. Synthetic population generation without a sample. Transportation Science. 2013;47(2):266–79.

17. Müller K, Axhausen KW, Axhausen KW, Axhausen KW. Preparing the Swiss Public-Use Sample for generating a synthetic population of Switzerland: Eidgenössische Technische HochschuleZürich, IVT, Institute for Transport Planning and Systems; 2012.

18. Zhu Y, Ferreira J. Synthetic population generation at disaggregated spatial scales for land use and transportation microsimulation. Transportation Research Record: Journal of the Transportation Research Board. 2014 (2429):168–77.

19. Harland K, Heppenstall A, Smith D, Birkin M. Creating realistic synthetic populations at varying spatial scales: a comparative critique of population synthesis techniques. Journal of Artificial Societies and Social Simulation. 2012;15(1):1–15.

20. Ryan J, Maoh H, Kanaroglou P. Population synthesis: Comparing the major techniques using a small, complete population of firms. Geographical Analysis. 2009;41(2):181–203.

21. Lomax N, Norman P. Estimating population attribute values in a table:”get me started in” iterative proportional fitting. The Professional Geographer. 2016;68(3):451–61.

22. Simpson L, Tranmer M. Combining Sample and Census Data in Small Area Estimates: Iterative Proportional Fitting with Standard Software. The professional Geographer. 2005;57(2):222–34.

23. Ma L, Srinivasan S. Synthetic Population Generation with Multilevel Controls: A Fitness-Based Synthesis Approach and Validations Computer-Aided Civil and Infrastructure Engineering 2015;30:135–50.

24. Williamson P, Birkin M, Rees P. The estimation of population microdata by using data from small area statistics and samples of anonymised records. Environment and Planning A abstract. 1998;30(5):785–816.

25. Voas D, Williamson P. An evaluation of the combinatorial optimisation approach to the creation of synthetic microdata. Population, Space and Place. 2000;6(5):349–66.

26. Price KV, Storn RM, Lampinen JA. Differential evolution: a practical approach to global optimization. Berlin; New York: Springer; 2005. xix, 538 p. p.

27. Gretton A, Borgwardt KM, Rasch MJ, Schölkopf B, Smola A. A kernel two-sample test. Journal of Machine Learning Research. 2012;13(Mar):723–73.

28. Fowler EE, Lu B, Heine JJ. A comparison of calibration data from full field digital mammography units for breast density measurements. Biomedical engineering online. 2013;12:114. PubMed PMID: 24207013. Pubmed Central PMCID: 3829208.

29. Heine JJ, Cao K, Rollison DE. Calibrated measures for breast density estimation. Acad Radiol. 2011 May;18(5):547–55. PubMed PMID: 21371912. Epub 2011/03/05.eng.

30. Heine JJ, Cao K, Rollison DE, Tiffenberg G, Thomas JA. A Quantitative Description of the Percentage of Breast Density Measurement Using Full-field Digital Mammography. Acad Radiol. 2011 May;18(5):556–64. PubMed PMID: 21474058. Epub 2011/04/09. eng.

31. Heine JJ, Fowler EEE, Flowers CI. A comparison of calibrated and non-calibrated breast density measurements with full field digital mammography Acad Radiol. 2011;18:1430–6.

32. Cacoullos T. Estimation of a multivariate density. Annals of the Institute of Statistical Mathematics. 1966;18(1):179–89.

33. Gramacki A. Nonparametric kernel density estimation and its computational aspects. Cham, Switzerland: Springer International Publishing AG; 2018.

34. Gramacki A, Gramacki J. FFT-Based Bandwidth Selector for Multivariate Kernel Density Estimation 2016 12 May, 2016;arXiv:1511.07482v3 [Stat.CO].

35. Duong T, Hazelton M, L. Cross-validation Bandwidth Matrices for Multivariate Kernel Density Estimation. Scandinavian Journal of Statistics. 2005;32:485–506.

36. Morel P. Gramm: grammar of graphics plotting in Matlab. Journal of Open Source Software. 2018;3(23).

37. Vernizzi G, Nakai M. A Geometrical Framework for Covariance Matrices of Continuous and Categorical Variables. Sociol Method Res. 2015 Feb;44(1):48–79. PubMed PMID: WOS:000350160200002. English.

